# Tonically active neurons in the monkey dorsal striatum signal outcome feedback during trial-and-error search behavior

**DOI:** 10.1101/2020.07.17.209601

**Authors:** Hitoshi Inokawa, Naoyuki Matsumoto, Minoru Kimura, Hiroshi Yamada

## Abstract

An animal’s choice behavior is shaped by the outcome feedback from selected actions in a trial-and-error approach. Tonically active neurons (TANs), presumed cholinergic interneurons in the striatum, are thought to be involved in the learning and performance of reward-directed behaviors, but it remains unclear how TANs are involved in shaping reward-directed choice behaviors based on the outcome feedback. To this end, we recorded activity of TANs from the dorsal striatum of two macaque monkeys (*Macaca fuscata*; 1 male, 1 female) while they performed a multi-step choice task to obtain multiple rewards. In this task, the monkeys first searched for a rewarding target from among three alternatives in a trial-and-error manner and then earned additional rewards by repeatedly choosing the rewarded target. We found that a considerable proportion of TANs selectively responded to either the reward or the no-reward outcome feedback during the trial-and-error search, but these feedback responses were not observed during repeat trials. Moreover, the feedback responses of TANs were similarly observed in any search trials, without distinctions regarding the predicted probability of rewards and the location of chosen targets. Unambiguously, TANs detected reward and no-reward feedback specifically when the monkeys performed trial-and-error searches, in which the monkeys were learning the value of the targets and adjusting their subsequent choice behavior based on the reward and no-reward feedback. These results suggest that striatal cholinergic interneurons signal outcome feedback specifically during search behavior, in circumstances where the choice outcomes cannot be predicted with certainty by the animals.

**Highlights:**

- TANs signal reward and no-reward outcome feedback when monkeys made search behaviors
- TANs respond regardless of predicted reward probability or chosen target location
- TANs may signal feedback outcomes when rewards cannot be predicted with certainty

## Introduction

Animals’ choice behaviors are shaped by the outcome feedback of their actions during trial-and-error searches. The striatum, an input stage of the basal ganglia, is involved in the learning and performance of reward-directed behaviors (Hikosaka et al., 2000; Schultz et al., 2003; Balleine et al., 2007) under the influence of two dominant neuromodulators: dopamine and acetylcholine. Dopamine neurons convey reward prediction error signals to the striatum, allowing animals to learn reward-directed behaviors (Montague et al., 1996; Schultz et al., 1997; Satoh et al., 2003; Kawagoe et al., 2004; Morris et al., 2004). Cholinergic signals originate from cholinergic interneurons in the striatum (Kawaguchi et al., 1995), are controlled by thalamic inputs (Matsumoto et al., 2001; Yamanaka et al., 2018), and interact with the dopamine signal (Cragg, 2006; Calabresi and Di Filippo, 2008; Surmeier et al., 2011). Although tonically active neurons (TANs, presumed cholinergic interneurons) signal the motivational outcomes of actions (Ravel et al., 2003; Morris et al., 2004; Yamada et al., 2004), it remains unclear how TANs signal outcome feedback for shaping reward-directed choice behavior in the striatal circuit.

In the striatum of behaving animals, medium spiny projection neurons, cholinergic interneurons, and parvalbumin-containing GABAergic interneurons are electrophysiologically characterized as phasically active neurons (PANs), TANs, and fast-spiking neurons (FSNs), respectively (Inokawa et al., 2010; Yamada et al., 2016). The activities of PANs (output neurons) and FSNs (inhibitory interneurons) are observed in relation to the learning and performance of reward-directed behaviors (Kawagoe et al., 1998; Jog et al., 1999; Schultz et al., 2003; Samejima et al., 2005; Balleine et al., 2007; Yamada et al., 2007; Lau and Glimcher, 2008; Schmitzer-Torbert and Redish, 2008; Gage et al., 2010; Lansink et al., 2010; Cai et al., 2011; Garenne et al., 2011; Yamada et al., 2016). Cholinergic signals modulate the activity of these striatal neurons (Calabresi et al., 2000; Goldberg et al., 2012; Schulz and Reynolds, 2013). TANs respond to behavioral outcomes and their predictive stimuli (Ravel et al., 2003; Morris et al., 2004; Yamada et al., 2004), and these responses seem to be strongly influenced by thalamic activities, which occur at unexpected environmental change that requires animals to adapt (Yamanaka et al., 2018). Moreover, responses of TANs are established throughout learning (Aosaki et al., 1994b; Blazquez et al., 2002). Thus, TANs seemed to be involved in shaping the choice behavior based on the outcome feedback.

The responses of TANs have been recently examined in animals during various cognitive and motor performances of reward-based decision-making tasks (Atallah et al., 2014; Franklin and Frank, 2015; Nougaret and Ravel, 2015; Stalnaker et al., 2016; Xiao et al., 2018). Among these, many potential roles of TANs in signaling outcome feedback have been suggested: TANs provide signals regarding reward prediction errors that are specific for learning stimulus-reward associations (Apicella et al., 2011), temporal timing for learning that are coincident with dopamine reward prediction error signals (Morris et al., 2004; Joshua et al., 2008), the identification of less attractive situations for animals (Nougaret and Ravel, 2015), the levels of uncertainty in stochastic environments (Franklin and Frank, 2015), switches in reinforcement contingencies (Atallah et al., 2014), and/or the belief state for associative learning (Xiao et al., 2018). Because these potential possibilities exist, we lack a common understanding of how TANs are involved in shaping the reward-directed choice behavior. Thus, it is worth elucidating the role TANs play in signaling the outcome feedback for shaping reward-directed choices during trial-and-error behaviors.

Herein, we examined activity of TANs using a reward-based decision-making task, called the multi-step choice task, during which monkeys searched for a rewarding target based on feedback, and obtained additional rewards by repeating the rewarded choices. We found that TANs detected reward and no-reward feedback outcomes specifically when monkeys performed trial-and-error searches, supporting the notion that TANs signal outcome feedback when the choice outcomes are not predicted with certainty by the animals, i.e., when animals needed to learn from the behavioral consequences.

### Experimental Procedures

All details of the behavioral task and the monkeys’ task performances have been described previously (Yamada et al., 2011). We have also described the activity of PANs and FSNs in our previous reports (Muranishi et al., 2011; Yamada et al., 2011; Yamada et al., 2013; Yamada et al., 2016); however, the TAN activity has not been reported previously, except for the spike waveforms for cell type identifications (Yamada et al., 2016).

### Experimental animals

We used two Japanese monkeys (*Macaca fuscata*: monkey RO, male, 9.4 kg; monkey TN, female, 6.3 kg). All experimental procedures were approved by the Animal Care and Use Committee of Kyoto Prefectural University of Medicine and conducted in compliance with the US Public Health Service’s Guide for the Care and Use of Laboratory Animals. All surgeries were performed under sterile conditions. Anesthesia was induced with ketamine hydrochloride (10 mg/kg, intramuscularly), followed by sodium pentobarbital (Nembutal, 27.5 mg/kg, intraperitoneally). Ketamine hydrochloride was additionally administrated when necessary. Heart rates and respiration were carefully monitored. Four head-restraining bolts and one stainless-steel recording chamber were implanted on the skull of each monkey under stereotaxic guidance. The recording chamber was laterally positioned at a 45° angle.

### Multi-step choice task

The monkeys performed a choice task to obtain multiple rewards through a series of choices (Fig. 1). In the task, monkeys searched for a rewarding target among multiple alternatives by adjusting their choices, i.e., shifting their choices after no-reward feedback, while repeating the rewarded choice after reward feedback.

**Figure 1.**
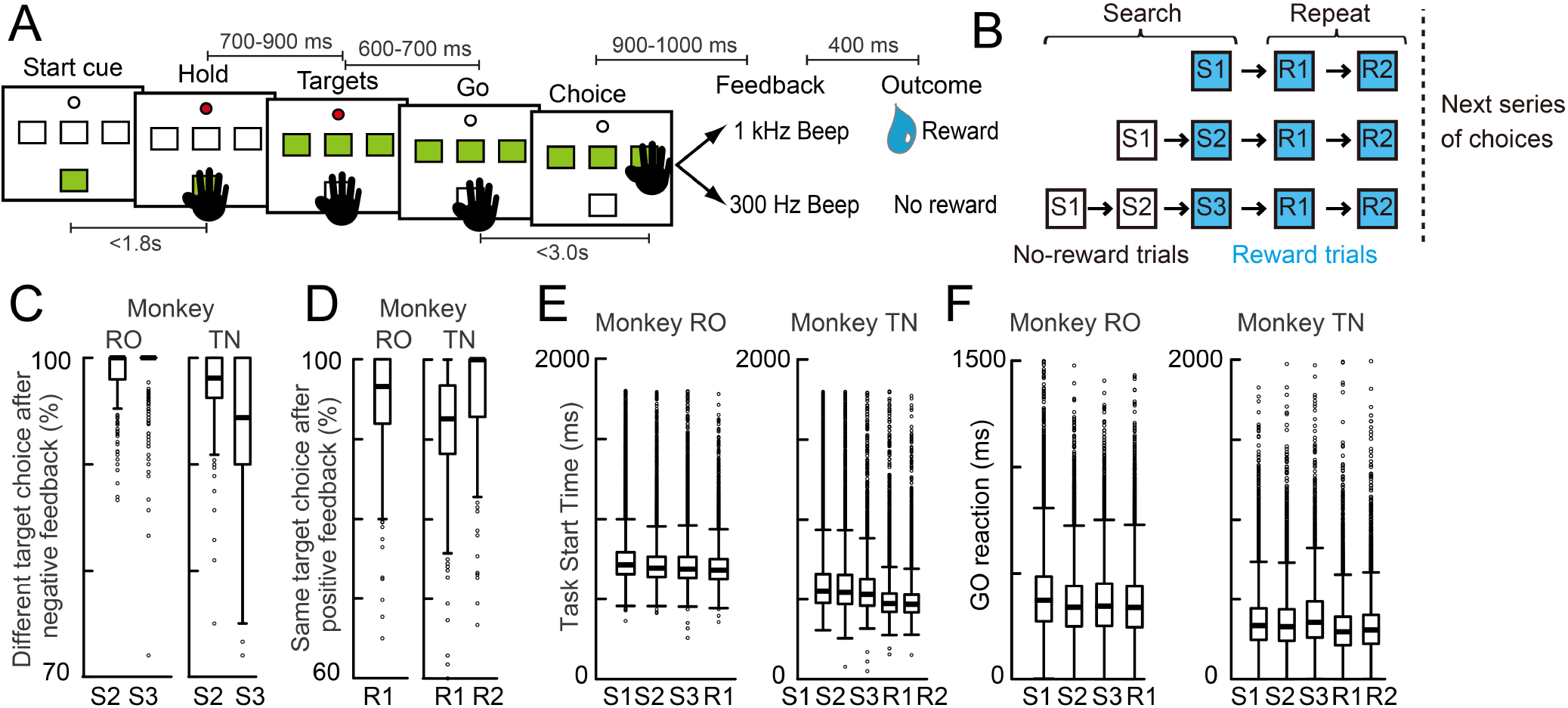
Multi-step choice task. ***A***, Schematic drawing of the behavioral events in a single trial. ***B***, Schematic drawing of a series of trials to obtain multiple rewards. Monkeys find a rewarding target in the first (S1), second (S2), or third (S3) search trial and obtain additional rewards in the subsequent repeat trials (R1 and R2). Monkeys obtained rewards twice (RO, no R2 trials) or three times (TN) through a series of choices. ***C***, Percentages for choosing a different target (search choice) in the S2 and S3 trials from the last no-reward choice in the S1 and S2 trials, respectively. ***D***, Percentages for choosing the same target (repeat choice) in the R1 and R2 trials from the last rewarded choice, respectively. ***E***, Task start times (latency from the onset of the start LED to the depression of the start button) in each trial type. ***F***, Choice reaction times (latency from the GO signal to the release of the start button) for each trial type. Graphs ***C-E*** have been previously shown in Yamada et al. (2011).

The monkeys were trained to sit on a primate chair facing a small wood panel placed 20.5 cm in front of their faces. Four buttons with light-emitting diodes (LEDs) and one small LED were embedded on the panel, as follows: a small rectangular start button with a green LED (start button, 14 × 14 mm) at the bottom, three target buttons with green LEDs (target buttons, 14 × 14 mm) in the middle row, and a small red LED (go LED, 9 mm in diameter) just above the center target button (Fig. 1A). After illumination of the start button (start cue), the monkeys depressed the start button with their right hand, contralateral to the neuronal recording. Following this, the go LED was turned on for 700–900 ms and the three target buttons were illuminated. The monkeys depressed the start button for 600–700 ms until the go LED was turned off (GO signal), after which they released the start button and depressed one of the three illuminated target buttons within 3 s. If the reward target button was depressed, a high-tone beep (1000 Hz for 200 ms, positive feedback) sounded following a delay of 900-1000 ms, and reward water (0.032 mL/kg weight) was delivered 400 ms after the disappearance of the beep sound. If a no-reward target button was depressed, a low-tone beep (300 Hz for 200 ms, negative feedback) sounded, and no reward was administered (Fig. 1A). The start LED was illuminated after a 5-s delay (inter-trial interval, ITI).

The monkeys searched for a rewarding target among the three target buttons based on the no-reward outcome feedback in the first (S1), second (S2), or third (S3) trial during search trials (Fig. 1B). Once the monkeys found a rewarding target, they obtained additional rewards in subsequent (R1 and R2) trials by reselecting the rewarded target. Monkeys obtained the rewards twice (RO) or three times (TN) through a series of choices. All four green LEDs were simultaneously flashed for 1.0 s at 2.0 s after the monkeys released the target button during the final repeat trial, which indicated that a single series of choices had ended. The next series of trials began 5 s after the flash of all buttons, with a new reward target at a random location.

### Single neuron recording in the striatum

Single neuron activity of TANs was recorded from the dorsal striatum, i.e., caudate nucleus and putamen, between A14 and A28. We used epoxy-coated tungsten microelectrodes and a template-matching algorithm to isolate single neuron discharges. We differentiated TANs from FSNs and PANs by their background discharge rates and spike waveforms (Yamada et al., 2016). Discharges of single neurons were recorded during approximately 170 and 200 trials in monkey RO and TN, respectively. This recording session was composed of 50-60 series of choices (a series of choices included 2-4 and 3-5 trials in monkey RO and TN, respectively, with a new reward target at a random location in each series of choices).

### Histological reconstruction

After all experiments were performed, small electrolytic lesions were made in the caudate nucleus and putamen by passing direct anodal current (20 μA) through tungsten microelectrodes for 30 s in each location. The monkeys were deeply anesthetized with an overdose of pentobarbital sodium (90 mg/kg, intravenously) and perfused with 4% paraformaldehyde in 0.1 M phosphate buffer through the left ventricle. Coronal sections of the striatum, 50 μm in thickness, were placed on slides and stained with cresyl violet. The tracks of the microelectrode through the striatum were reconstructed on the histology sections using the electrolytic lesion marks as reference points.

### Data analysis

#### Analysis of behavioral performance

We reused previously reported behavioral data during the multi-step choice task to support this current study (Yamada et al., 2011). Behavioral data were obtained from a total of 98,604 and 61,299 trials in 583 and 303 recording sessions in monkeys RO and TN, respectively. The percentage of finding a rewarding target in each trial type (S1, S2, S3, R1, and R2) was determined as the ratio of the number of rewarded trials to the total number of trials during each recording session. Task start times (from the illumination of the start LED to the depression of the start button) were compared among trial types using multiple two sample comparisons corrected by the Bonferroni method to control for the family-wise significance level. Choice reaction times (time from the GO signal to release the start button) were compared among trial types using multiple two sample comparisons after applying the Bonferroni correction method. The percentage of same or different target choices was estimated in each trial type after reward and no-reward feedback, respectively.

#### Identification of TAN responses to the start cue and feedback beeps

For each TAN, histograms of the neuronal discharge rates were constructed before and after particular behavioral events. A significant increase or decrease in the discharge rate was determined by comparing the discharge rate during an 80 ms test window with the rate during the 400 ms baseline period just before the onset of the start LED (Wilcoxon two-sample test, statistical threshold: *p* < 0.05). We detected significant changes in the discharge rate after each of the five behavioral events, i.e., the start cue during search trials, the start cue during repeat trials, the reward feedback during search trials, the no-reward feedback during search trials, and the reward feedback during repeat trials with Bonferroni correction (i.e., significance is accepted as *p* < 0.01) (Morris et al., 2004). In each of the five behavioral events, comparisons were made by shifting the 80 ms test window in 10 ms steps during the 600 ms period following the event. The activity was regarded as significantly different from the baseline if more than five consecutive comparisons between the test and baseline window reached statistical significance of *p* < 0.01. The test window was further shifted until the discharge rate within the test window was not significantly different from the baseline firing rate for five consecutive steps. Onset of the significant activity change was defined as the beginning of the window that occurred the earliest among the first five consecutive steps. Disappearance of significant activity change was defined as the beginning of the earliest window among the last five consecutive steps.

The significant activity changes were regarded as TAN responses in each of the five behavioral events. In addition, we also detected the responses of TANs to other behavioral events, such as start button push, target onset, GO signal, target button push, and onset of reward during search and repeat trials. We used the same statistical significance level as that used for the detection above (i.e., significance is accepted as *p* < 0.01) to avoid too strong correction. Percentages of the TAN responses during search and repeat trials were compared using chi-squared tests with a statistical threshold of *p* < 0.05 or binomial tests with a chance level of 5%. The magnitudes of TAN responses were compared using two-sample *t*-tests or ANOVAs with a statistical threshold of *p* < 0.05.

#### Estimation of the mean response magnitudes of TAN

The increase and decrease in the discharge rates of TAN responses were estimated for each neuron using methods similar to those reported by Morris et al. (2004). Histograms of the neuronal discharge rates were constructed using 1 ms bins with a Gaussian kernel (σ = 25 ms), and the baseline firing rates were subtracted from the mean firing rate during the task period. The increase and decrease from baseline firing rates were estimated separately during the 600 ms task period.

#### Identification of the temporal patterns of TAN responses

The temporal patterns of TAN responses were classified using principal component analysis (PCA) followed by the k-means clustering algorithm. PCA was applied once to the peri-stimulus time histogram (PSTH) data during 400 ms period after reward and no-reward feedback and those after start cue during search and repeat trials. The k-means algorithm was used to classify four types of responses according to the PC1 to PC3 scores, since TANs usually showed some of the four components of the responses: short latency facilitation < 250 ms (i.e., initial facilitation), suppression at around the 200 ms, facilitation following the suppression and suppression following the facilitation (Yamada et al., 2004). The four response types were accordingly labeled as initial facilitation (IF), suppression only (S), suppression followed by facilitation (SF), and facilitation only (F). The non-responsive TANs were excluded from the analysis. The proportions of these four response types were compared using chi-squared tests, with the statistical threshold set at *p* < 0.05.

## Results

### Monkey’s Behavior

During the multi-step choice task, the two monkeys chose a target button with their hand (Fig. 1A) and searched for a rewarding target from among the three target buttons based on no-reward outcome feedback (Fig. 1B, search trials). After finding a rewarding target, they repeated the choice of the rewarded target to earn additional rewards (Fig. 1B, repeat trials). The monkeys received two (monkey RO) or three (monkey TN) times of rewards through a series of choices. Although details of the behavioral training and performances of these two monkeys in the multi-step choice task have previously been reported (Yamada et al., 2011), they have been rereported here to support this present study.

The two monkeys learned how to search for a reward target by pressing one of three target buttons in each trial (Fig. 1A). After completing the training, the percentages of finding a rewarding target gradually increased over the first (S1, mean ± S.D. 33.0 ± 3.5% and 32.3 ± 3.0% in monkey RO and TN, respectively), second (S2, 50.0 ± 4.5% and 48.9 ± 4.8% in monkey RO and TN, respectively), and third (S3, 88.7± 9.0 % and 81.6 ± 10.8 % in monkey RO and TN, respectively) trials during a series of choices (Fig. 1B). The percentages to find a reward in the S3 trials was not 100% due to all 3 target options being presented, thus, the monkeys sometimes chose one of the non-rewarded targets. During the search trials, monkeys shifted their choices following no-reward feedbacks in more than 90% of the trials (Fig. 1C). Once the monkeys found a reward target, they obtained one or two additional rewards by repeating the same target choice in an additional one (R1, Monkey RO) or two trials (R1 and R2, Monkey TN). The monkeys repeated the rewarded choice after receiving reward feedback in about 95% of the trials (Fig. 1D, Monkey RO, R1 96.1 ± 4.7%, Monkey TN, R1, 93.2 ± 6.7%, R2, 95.6 ± 6.0%).

Behavioral measures were dependent on the search and repeat trials, as well as on the reward probability during a series of trials (Fig. 1E). The task start time (time from the illumination of the start LED to the depression of the start button) decreased with the increase in the percentage to find a reward (Fig. 1E, Bonferroni correction, *p* < 0.001, except between the S2 and S3 trials of monkey RO, *p* = 0.99). The task start times were shorter in the repeat trials than in the search trials (*p* < 0.001 in all cases). The reaction time to choose a target button was also dependent on the search and repeat trials (Fig. 1F). These choice reaction times were longer in the search trials as compared to repeat trials (Bonferroni correction, *p* < 0.001, except between the S2 and R1 trials of monkey RO, *p* = 0.079). They were the longest in S3 trials in monkey TN. In monkey RO, the reaction times were longer in S3 trials compared to S2 trials. This might be because the ability to memorize the location of unrewarded target is higher in monkey RO than that in monkey TN (S3, 88.7± 9.0 % and 81.6 ± 10.8 % in monkey RO and TN, respectively). Thus, monkeys adjusted their behavior after reward and no-reward feedbacks during the trial-and-error searches and the following repeat trials through a series of choices.

### Feedback response of TANs during the multi-step choice task

Previously, we reported the activity of 292 PANs and 31 FSNs (Yamada et al., 2011; Yamada et al., 2013; Yamada et al., 2016) recorded from the striatum of the same two monkeys during the performance of multi-step choice task. In the present study, we focused on analyzing the activity recorded from 149 TANs in the striatum (Fig. 2, 92 and 57 TANs in monkey RO and TN, respectively).

**Figure 2.**
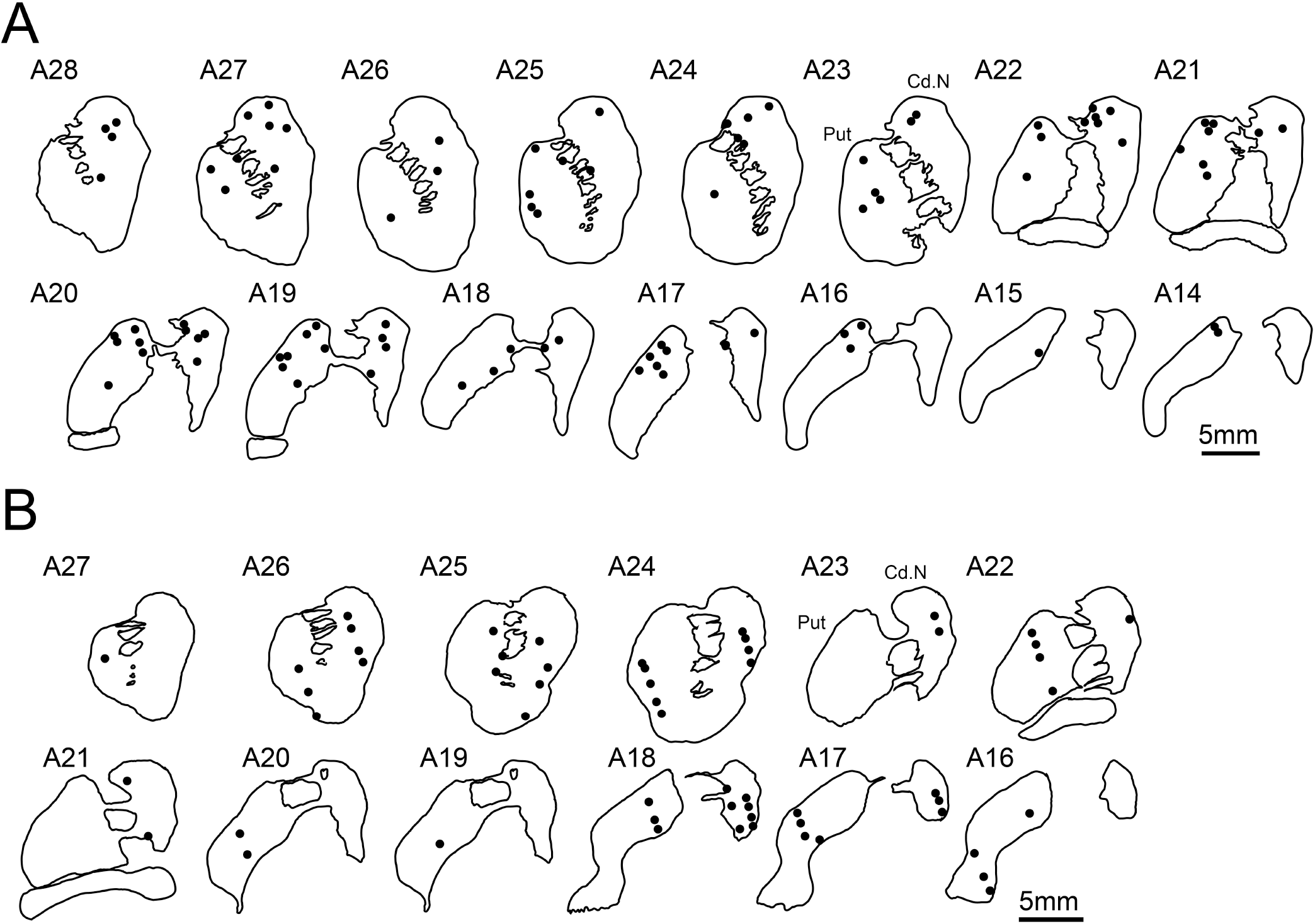
Recording location of TANs in the striatum. ***A,*** Locations of the recorded TANs in monkey RO. ***B,*** Same as ***A*** but for the recorded TANs in monkey TN. The anterior commissure is located at the A22-20 level. Cd.N, caudate nucleus, Put, putamen.

An example of TAN activity recorded during the performance of multi-step choice task after the occurrence of reward and no-reward feedback beeps is shown in Fig. 3A. After the monkey received no-reward feedback during search trials (Fig. 3A, S1 and S2, green), TAN activity decreased, followed by an increase in the discharges. In this example, the TAN also responded to reward feedback when the monkeys searched for a rewarding target irrespective of the trial type (S1, S2, and S3 trials) (Fig. 3A, blue). However, significant responses were not observed once the monkey found the rewarding targets and repeated the rewarded choices (Fig. 3A, magenta, R1). The responses of this example of TAN were evoked only during search trials (Fig. 3B, green and blue) and not during repeat trials (Fig. 3B, magenta). A further example in which TAN showed clear decrease responses to reward feedback beeps during search trials is shown in Fig. 3C and D (blue). Only slight increases in activity were shown after reward feedbacks in the first repeat trials (Fig. 3D, magenta, R1).

**Figure 3.**
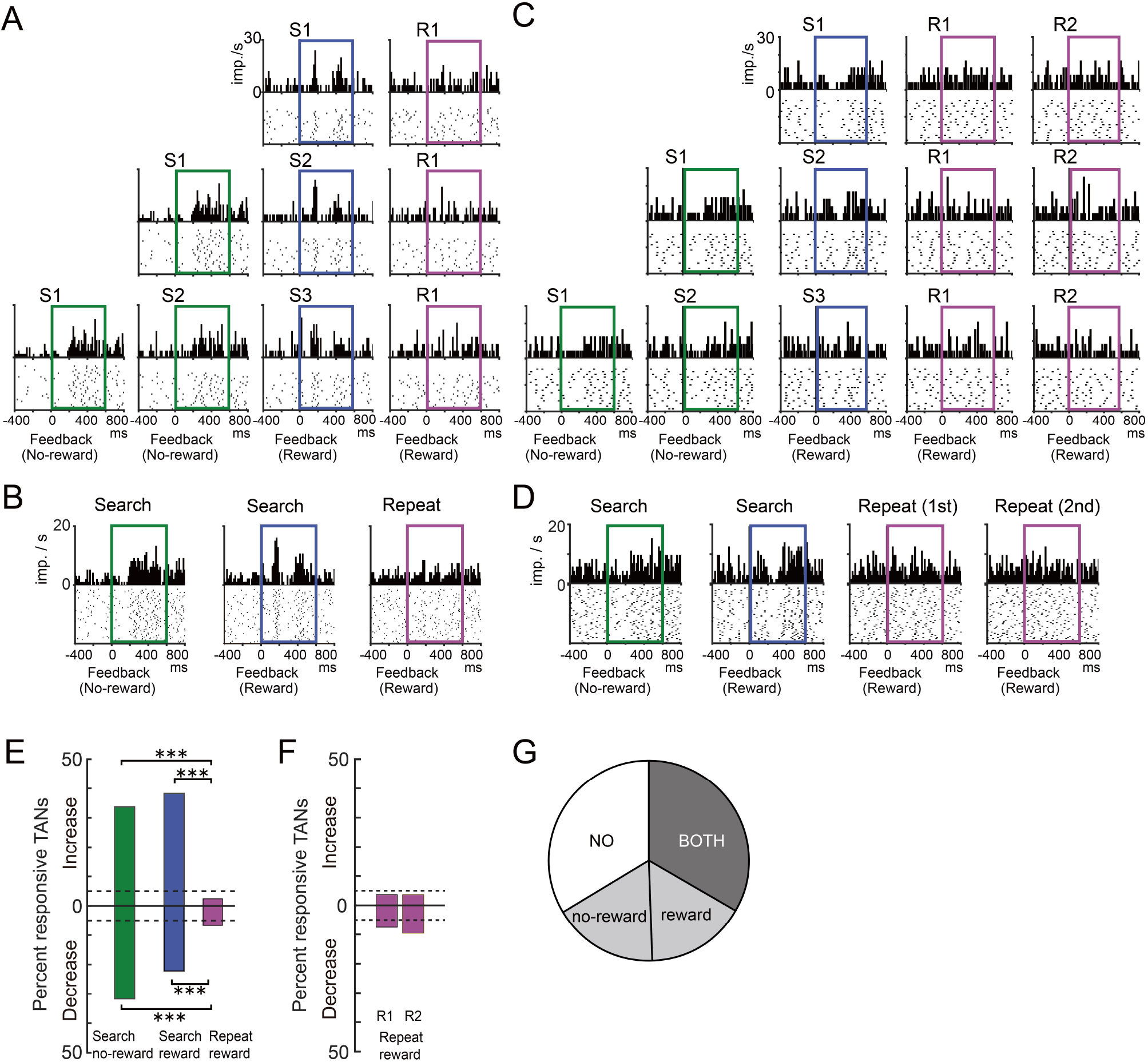
Search trial-specific feedback responses of TANs. ***A***, Peri-stimulus time histograms (15 ms bins) and raster plots are depicted for example TAN responses after reward (blue and magenta) and no-reward (green) feedback in each trial type (S1, S2, S3, and R1). Colored squares indicate the time window to detect the responses of the TAN. The TAN response was recorded in monkey RO. ***B***, Responses of the same TAN shown in ***A***, but for the averaged histograms after no-reward feedback during search trials (green, S1 and S2), after reward feedback during search trials (blue, S1, S2, and S3), and after reward feedback during repeat trials (magenta, R1). Colored squares indicated the time window to detect the response of TANs. ***C-D,*** same as ***A-B***, but for another example TAN response in monkey TN (S1, S2, S3, R1, and R2 trial types). ***E***, Proportion of feedback responses of TANs showing significant activity increases and decreases. ***F***, Proportion of feedback responses of TANs during repeat trials in monkey TN. The responses were detected in first and second repeat trials (R1 and R2, respectively). In ***E*** and ***F***, Dashed lines indicate the 5% chance level. \*\*\**p* < 0.001. ***G***, Proportion of TANs exhibiting feedback responses during search trials: BOTH, TANs that responded to both reward and no-reward feedback; reward, TANs that responded to reward feedback only; no-reward, TANs that responded to no-reward feedback only; NO, TANs that responded to neither reward nor no-reward feedback. In ***A***-***D***, imp./s is impulse per second.

We analyzed the reward and no-reward feedback responses of TANs during the search and repeat trials quantitatively. Both increases and decreases in TAN responses were frequently observed during search trials compared to repeat trials (Fig. 3E; chi-squared test: search no-reward vs. repeat reward: increase, X^2^[1] = 46.6, *p* < 0.001; decrease X^2^[1] = 28.1, *p* < 0.001; search reward vs. repeat reward: increase, X^2^[1] = 37.9, *p* < 0.001; decrease X^2^[1] = 13.2, *p* < 0.001). Almost no response was observed during repeat trials at the 5% chance level (Fig. 3E, magenta; binomial test: increase, 2.9%, 4/149, *p* = 0.257; decrease, 6.7%, 10/149, *p* = 0.342). During search trials, about half of the TANs showed a change in their activity (either an increase or a decrease) after reward (50.3%, 75/149) and no-reward (50.3%, 75/149) feedback, respectively, but during repeat trials, only a small proportion of TANs showed either response after reward feedback (9.4%, 14/149; chi-squared test: X^2^[1] = 70.7, *p* < 0.001). If we analyzed responses of TANs to reward feedbacks during repeat trials by differentiating the first and second repeat trials (R1 and R2 trials in monkey TN), only a small proportion of TANs showed responses during R1 and R2 trials (Fig. 3F, chi-squared test: R1 vs. R2: increase, X^2^[1] = 0.342, *p* = 0.559; decrease X^2^[1] = 1.67, *p* = 0.197). During search trials, about one third of TANs showed responses after both reward and no-reward feedback (Fig. 3G; 34.2%, 51/149), whereas another one third of TANs showed responses after either reward or no-reward feedback (32.2%; reward only, 16.1%, 24/149; no-reward only, 16.1%, 24/149). Thus, feedback responses of TANs were specifically observed when monkeys searched for a rewarding target, with exhibition of distinct responses between reward and no-reward feedbacks in considerable proportions.

The feedback responses of TANs showed distinct temporal patterns. For example, activity decreases were followed by increases after no-reward feedback (Fig. 3B, green), while after reward feedback, short-latency activity increases were followed by decreases and then subsequent activity increases (Fig. 3B, blue). We classified the temporal patterns of responses into the S, SF, F, and IF types by using PCA followed by k-means clustering algorithm (see Materials and Methods section for details; Fig. 4A-C). During search trials, all four types of responses were observed after both reward and no-reward feedback (Fig. 4A and B), but they were different in proportion (Fig. 4D). After no-reward feedback, the S, SF, and F types were observed, while the IF type was rarely observed (Fig. 4D, left). In contrast, after reward feedback, a considerable number of IF type responses were observed (Fig. 4D, right), while the F type was rarely observed. The proportions of the four types of responses were significantly different between reward and no-reward feedback during the search trials (chi-squared test: X^2^[3] = 34.8, *p* < 0.001). An activity heat map of all recorded TANs (Fig. 4E) showed short-latency activity increases after reward feedback (middle column, red color after feedback); however, no such responses were observed after no-reward feedback (left column, blue color just after feedback). Thus, the temporal patterns of the responses dissociated the reward and no-reward feedback as a population. Note that a very small proportion of TANs responded to reward feedback during repeat trials (Fig. 4C, right column), however the magnitudes of the responses in the repeat trials were significantly smaller than those in the search trials (Fig. 4F; two sample *t*-test: increase, *t*[296] = 10.2, *p* < 0.001; decrease, *t*[296] = 6.59, *p* < 0.001).

**Figure 4.**
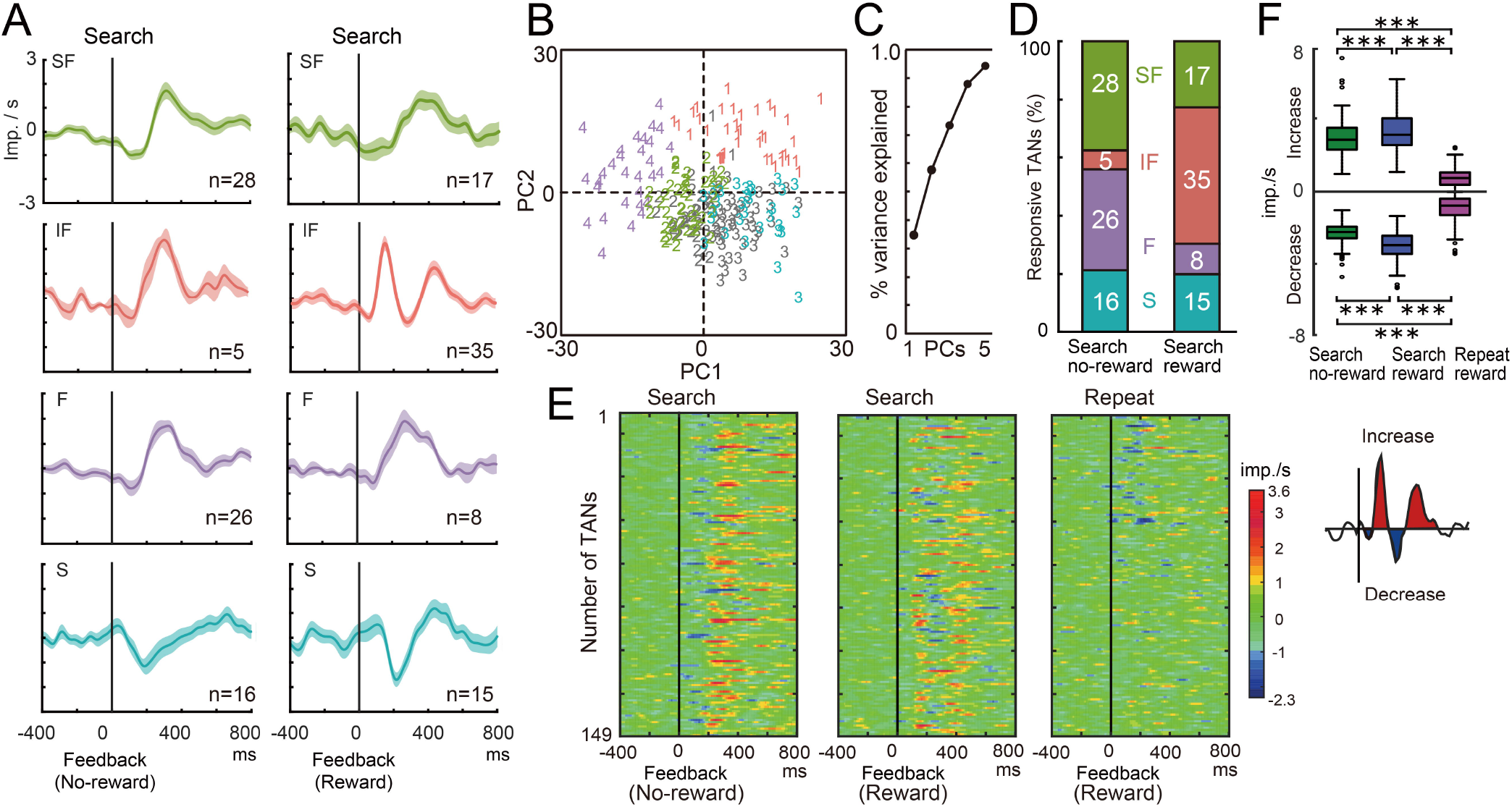
Temporal patterns of feedback responses differed between reward and no-reward feedback. ***A***, Population histograms of the four types of TANs after reward and no-reward feedback for search trials, including the suppression only (S), facilitation only (F), initial facilitation (IF), and suppression followed by facilitation (SF) types. ***B***, Score plot of the first two principal components (PC1 and PC2). Numerical numbers indicate the four clusters defined based on the k-means algorithm. Gray plots indicate the responses of TANs after start cue. **C,** Cumulative plot of the percent variance explained by the PCA shown against PC1 to PC5. ***D***, Proportions of the four types of TANs after reward and no-reward feedback during search trials. The number of TANs is shown in the bar histograms. ***E***, Heat map plots of the activity of all TANs after no-reward feedback during search trials (left), after reward feedback during search trials (middle), and after reward feedback during repeat trials (right). Each line represents neuronal discharges relative to baseline activity. In the three heat map plots, activity of the same neuron is arranged horizontally in the same row. ***F***, Mean response magnitudes of TANs after no-reward feedback during search trials (green), after reward feedback during search trials (blue), and after reward feedback during repeat trials (magenta). Inset below represents the TAN responses quantified as the increase and decrease from the baseline firing rate. \*\*\**p* < 0.001. In ***A***, ***E*** and ***F***, imp./s means impulse per second relative to the base line firing rates.

We then examined whether the search-specific feedback responses of TANs encoded the location of chosen targets using one-way ANOVAs. An example TAN showed no modulation of their responses by the location of the chosen target (Fig. 5A, one-way ANOVA, search reward: increase F[2, 44] = 0.131, *p* = 0.877 decrease F[2, 44] = 2.14, *p* = 0.129, search no-reward: increase F[2, 54] = 0.757, *p* = 0.474 decrease F[2, 54] = 2.30, *p* = 0.109). Of the 99 TANs that responded to either reward or no-reward feedback, only 8.1% (8/99) showed activity modulation by the location of the chosen target when the increase and decrease in activity were analyzed together (Fig. 4B; binomial test at 5% chance level, *p* = 0.162). Consistently, almost no target-dependent modulation was observed when we separately analyzed either increase or decrease in responses (increase, 3.0%, 3/99; decrease, 5.1%, 5/99). Thus, the search-specific responses of TANs did not signal the location of chosen targets.

**Figure 5.**
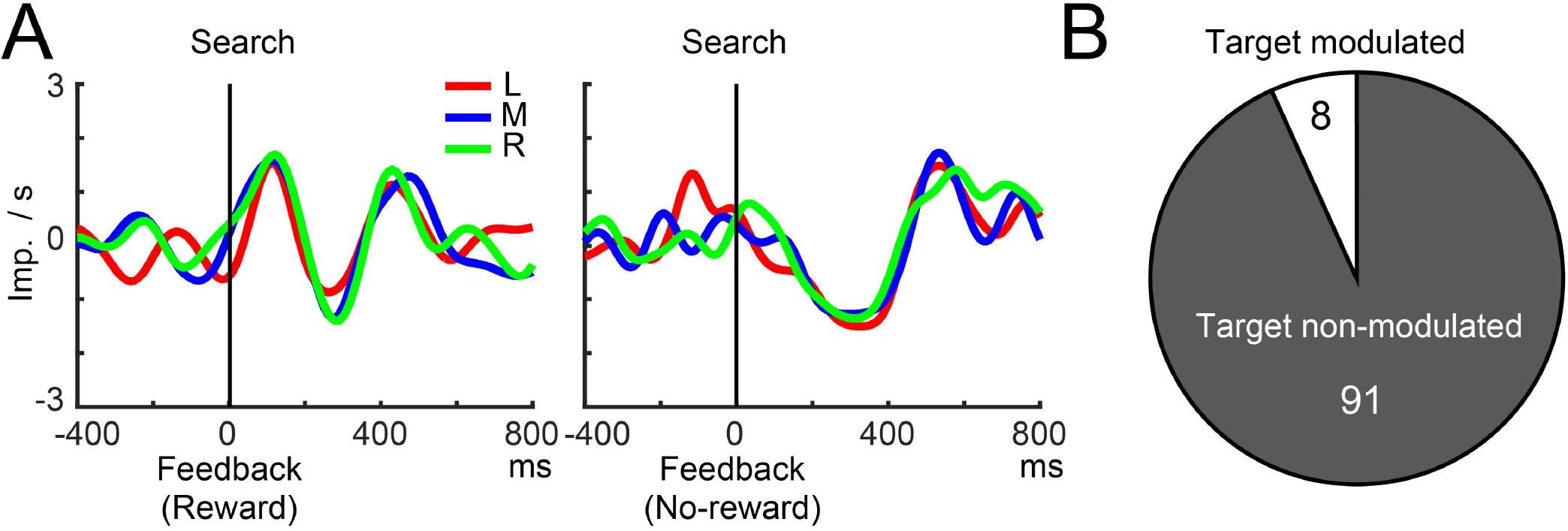
Feedback responses of TANs did not differ according to the location of chosen targets. ***A***, Peri-stimulus time histograms (15 ms bins) are depicted for example responses of TAN after reward and no-reward feedback during search trials. The histograms are shown for the different locations of chosen targets (L: left, M: middle, R: right). ***B***, Proportions of TANs modulated by the location of chosen targets. The number of TANs is shown in the pie-chart plot. imp./s means impulse per second relative to the base line firing rates.

We also examined whether the search-specific responses of TANs reflected the probability of finding a rewarding target, since reward prediction and its error, known as the reward-prediction-error signal, which is conveyed by dopamine neurons to the striatum (Enomoto et al., 2020), are critical for choice behavior. Neither increases nor decreases in activity after reward and no-reward feedback depended on reward probability as a population (Fig. 6A; search reward: one-way ANOVA, increase, F[2, 444] = 1.78, *p* = 0.17; decrease, F[2, 444] = 0.14, *p* = 0.881; search no-reward: increase, F[2, 418] = 2.60, *p* = 0.08; decrease, F[2, 418] = 2.65, *p* = 0.07). 6.1% (6/99) of TANs exhibited responses that were modulated by the reward probability (Fig. 6B; binomial test at 5% chance level, *p* = 0.64; increase, 6/99; decrease, 3/99). Moreover, responses of TANs after no-reward feedback were smaller than those after reward feedback (two-way ANOVA, increase, feedback type, F[1, 864] = 13.2, p < 0.001, decrease, feedback type, F[1,864] = 111.0, p < 0.001) consistent with the results mentioned above (Fig. 4F). Therefore, TANs encoded search-specific feedback signals without distinctions with respect to either the monkeys’ target choice or predicted reward probability, but with differentiation of reward and no-reward feedback in their response magnitudes. These results suggested that the feedback responses of TANs only discriminated whether the monkeys searched for a reward or not during a series of choices.

**Figure 6.**
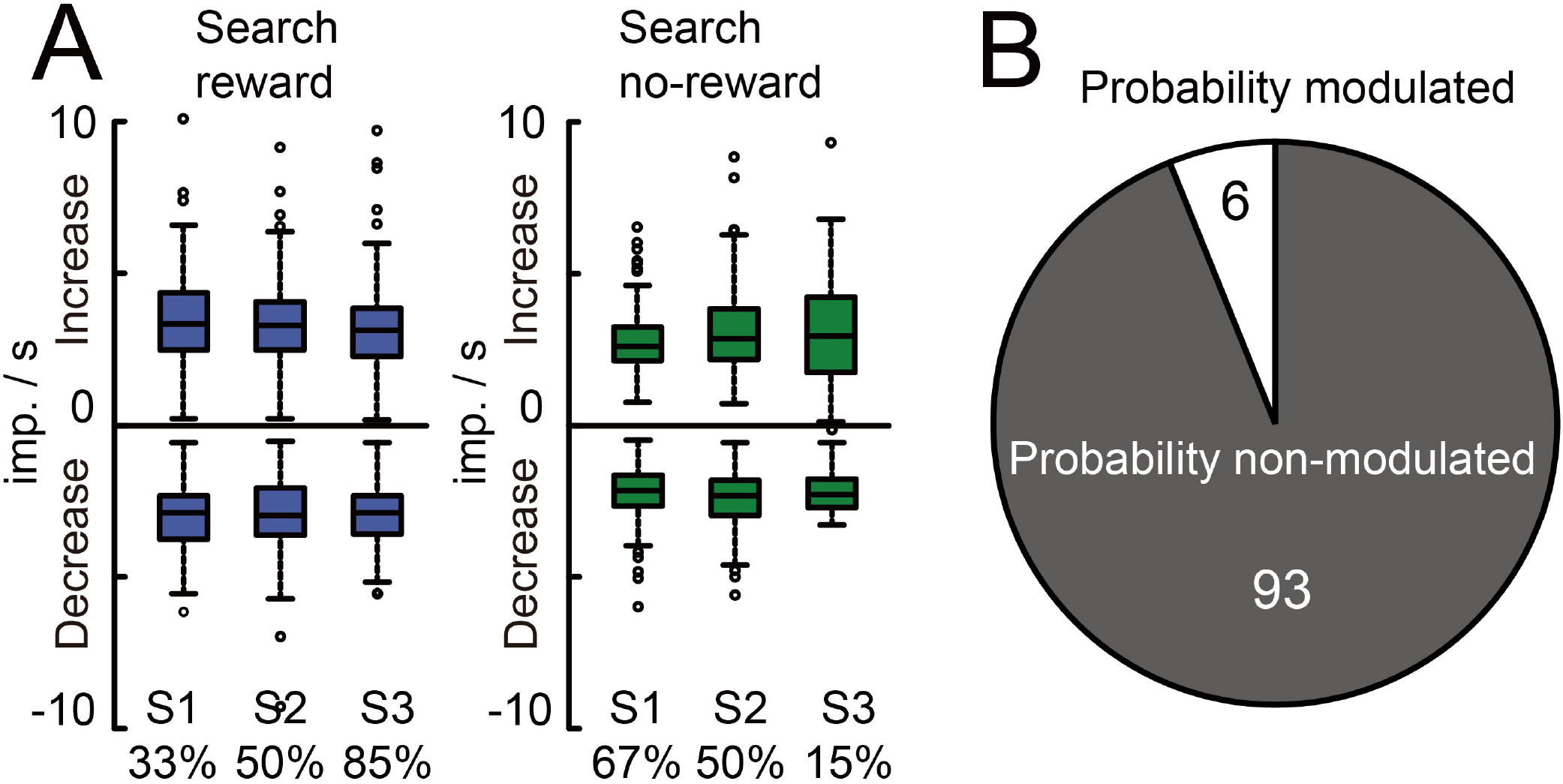
Feedback responses of TANs did not depend on the reward probability. ***A***, Mean response magnitudes after reward (blue) and no-reward (green) feedback are shown during search trials. Percentages below indicate the approximate probabilities of receiving or not receiving rewards in each trial type; imp./s is impulse per second relative to the base line firing rates. ***B***, Proportions of TANs modulated by the reward probability. The number of TANs is shown in the pie-chart plot.

### Search-specific feedback responses of TANs during errors

We examined whether search-specific feedback responses of TANs reflect monkey’s task performance to find a rewarding target. We analyzed the responses of TANs during S3 trials, in which monkeys made error choices about 15% of the trials. Since these error trials were not large in number, we analyzed aggregated population data of TANs. The population histogram of TANs to feedbacks in the S3 error trials were different from those of error trials in terms of the temporal pattern of the responses (Fig. 7A), consistent with the results as shown above (Fig. 4). The magnitude of the decreased responses was slightly smaller in no-reward feedback than in reward feedbacks in S3 trials (Fig. 7B. two sample *t*-test: increase, *t*[270] = 0.462, *p* = 0.644; decrease, *t*[270] = 2.69, *p* = 0.008), same as the weaker no-reward feedback responses in S1 and S2 trials (Fig. 6A). Thus, no-reward feedback responses of TANs when monkeys made error choices showed similar response properties to no-reward feedbacks when monkeys correctly chose un-chosen target buttons.

**Figure 7.**
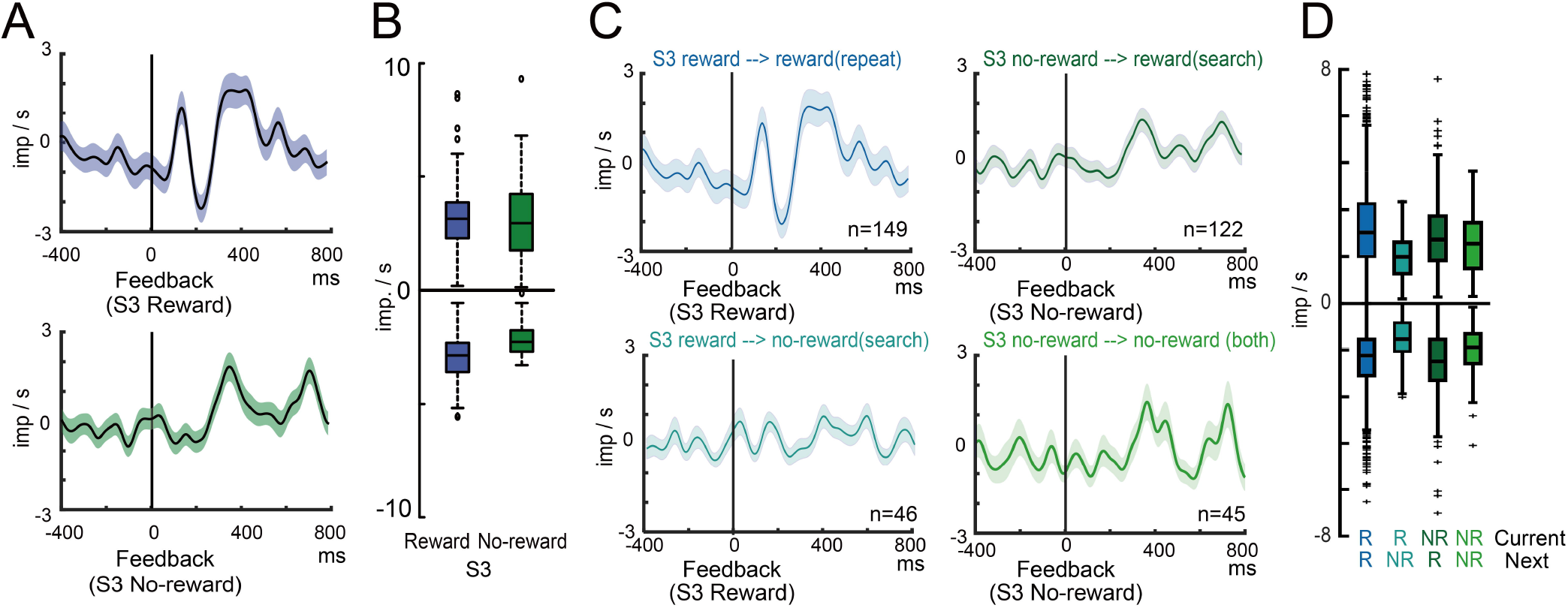
Feedback responses of TANs when monkeys made errors in S3 trials. **A,** Population average histogram of TANs to reward and no-reward feedbacks during S3 trials. **B,** Mean response magnitudes after reward (blue) and no-reward (green) feedback during S3 trials. **C,** Population average histogram of the TANs to reward and no-reward feedback during S3 trials for four different type of trials. Rewarded search choice then next reward choice, Rewarded search choice then next no-reward choice, no-reward search choice then next reward choice, and no-reward search choice then next no-reward choice. Number in each figure indicates number of TANs. **D,** Mean response magnitudes during S3 trials shown in C. R and NR indicate reward and no-reward, respectively. The TAN responses quantified as the increase and decrease from the baseline firing rate; imp./s means impulse per second relative to the base line firing rates.

To further examine whether responses of TANs reflect monkey’s task performances to find a rewarding target, we examined whether the feedback responses of TANs reflect the monkey’s choice behavior in the subsequent trials. We analyzed the population average responses of TANs during search trials by differentiating the monkey’s choice behavior in the next trials (Fig. 7C). Feedback responses of TANs to reward feedbacks were not clearly observed when monkeys made error choices in the next trials (Fig. 7C, left bottom). While the number of error choices in the next trials was not large, these differences were significant both in increase and decrease in their discharge rates (Fig. 7D, two-way ANOVA, increase: current feedback, F[1, 353] = 0.131, p = 0.657, next feedback, F[1, 353] = 21.7, p < 0.001, interaction, F[1, 353] = 24.2, p < 0.001, decrease: current feedback, F[1, 353] = 6.17, p = 0.013, next feedback, F[1, 353] = 34.3, p < 0.001, interaction, F[1, 353] = 7.49, p = 0.007). These results suggested that responses of TANs seemed to be related to monkey’s task performances to search a rewarding target.

### Feedback responses of TANs in the caudate nucleus and putamen

TANs in the caudate nucleus and putamen differentiate the cue stimulus associated with motivational outcomes (Yamada et al., 2004). We thus examined whether the search trial specific feedback responses of TANs are different between the caudate nucleus and putamen (67 and 82 TANs were recorded from the caudate nucleus and putamen, respectively). We found two significant differences of the TAN’s responses in the present study. The percentages of increased responses after no-reward feedbacks were higher in the putamen than in the caudate nucleus (search no-reward, increase, caudate, 16/67, 23.9%, putamen, 34/82, 41.5%, chi-squared test: X^2^[1] = 4.35, p = 0.037, decrease, caudate, 21/67, 31.3%, putamen, 26/82, 31.7%, X^2^[1] = 0, p = 0.999), while TANs did not show significant statistical differences after reward feedbacks (search reward, increase, caudate, 22/67, 32.8%, putamen, 35/82, 42.7 %., chi-squared test: X^2^[1] = 1.13, p = 0.289, decrease, caudate, 10/57, 14.9%, putamen, 23/82, 28 %., X^2^[1] = 0, p = 0.085). We also found that IF type response (cluster No.1) after reward feedback was predominant in the putamen compared to the caudate nucleus (Putamen, IF, 28, SF, 6, S, 11, F, 1, caudate, IF, 7, SF, 11, S, 4, F, 7, chi-squared test: X^2^[3] = 19.0, p-value < 0.001).

### Responses of TANs to start cues and other behavioral events

Lastly, we examined the responses of TANs to other behavioral events, such as the start cue, hold button push, onset of target, GO signal, choice, and reward delivery during search and repeat trials (Fig. 8A). TANs responded to the start cues during search and repeat trials. These responses were significantly frequent at the 5% chance level, especially for the decreased response (binominal test, increase, search, *p* = 0.012, repeat, *p* = 0.850; decrease, search, *p* < 0.001, repeat, *p* < 0.001). Population histogram of all TANs showed that they similarly responded to the start cues during search and repeat trials (Fig. 8B; one-way ANOVA: increase, F[2, 444] = 1.02, p = 0.36; decrease, F[2, 444] = 1.34, p = 0.26). Temporal patterns of these start cue responses were clearly different from those of feedback beeps (Fig. 8C-D), but proportions of the four types of responses were not significantly different between search and repeat trials (Fig. 8E, chi-squared test: X^2^[3] = 1.21, *p* = 0.55). Neither an increase nor decrease in activity after the onset of the start cue depended on the reward probability (one-way ANOVA: increase, F[3, 592] = 2.06, *p* = 0.10; decrease, F[3, 592] = 2.09, *p* = 0.10). Note that considerable proportion of TANs showed decrease responses significantly when monkeys pushed a target button (i.e., choice, Fig. 8A and F). These results indicated that the start cue responses of TANs did not dissociate behavioral context, search or repeat, in contrast to the feedback responses.

**Figure 8.**
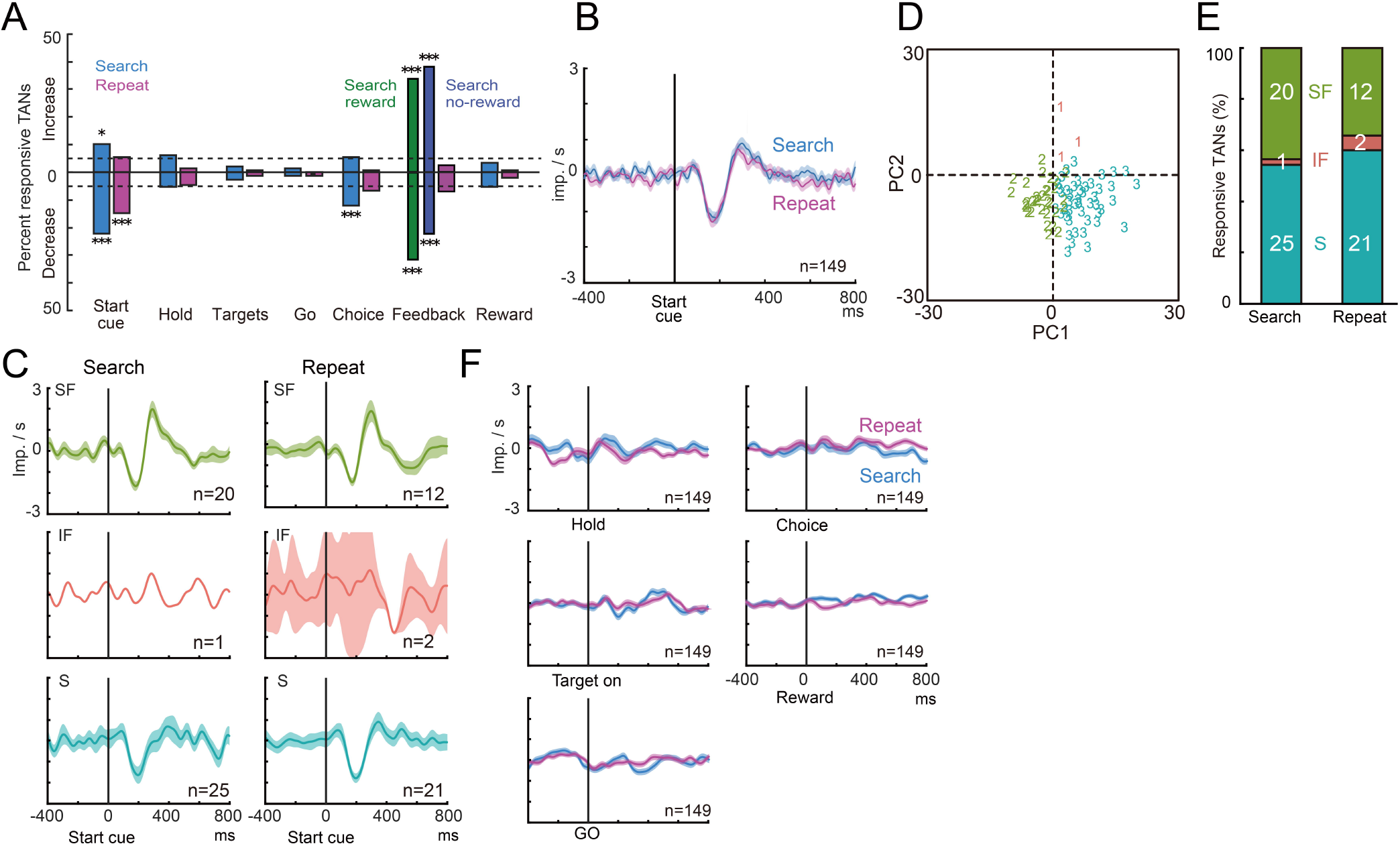
Responses of TANs to the start cue and other behavioral events. ***A***, Percentages of increase and decrease responses of TANs to all behavioral events during search and repeat trials. ***B***, Population histogram of all 149 TANs after start cue illumination during search and repeat trials. imp./s means impulse per second relative to the base line firing rates. Error bars indicate SEM. **C,** Population histograms of the three types of TANs after start cue, including the suppression only (S), initial facilitation (IF), and suppression followed by facilitation (SF) types. Facilitation only (F) was not observed. **D,** Score plot of the first two principal components (PC1 and PC2) to start cue responses during search and repeat trials. ***E***, Proportions of the four types of TANs after reward and no-reward feedback during search trials. F type was not observed. The number of TANs is shown in the bar histograms. **F,** Population histogram of all 149 TANs after holding start button (Hold), onset of target (Target on), GO signal (GO), target button push (Choice), and onset of reward (Reward). * and *** indicates *p* < 0.05 and *p* < 0.001, respectively.

## Discussion

Herein, we examined how TANs were involved in signaling outcome feedback during the multi-step choice task. We found that feedback responses of TANs were specifically observed when monkeys searched for a rewarding target using a trial-and-error approach; responses were not observed in repeat trials after the monkeys found a rewarding target. A considerable proportion of TANs responded to either or both feedback outcomes (reward and no-reward), with the responses exhibiting distinct temporal patterns between reward and no-reward feedback. Unambiguously, the search trial-specific feedback responses of TANs reflected neither the reward probability nor the location of chosen targets, indicating that TANs detected the outcome feedback when monkeys performed trial-and-error search behavior. These results suggest that striatal cholinergic interneurons signaled outcome feedback during trial-and-error searches, when monkeys learned from the feedback outcomes.

### Responses of TANs to outcome feedback and cue stimuli

We found that TANs showed responses to the outcome feedback and start cue during a series of choices to obtain multiple rewards. Our results were in partial agreement with previous studies in terms of the response modulation by reward probability (Morris et al., 2004; Apicella et al., 2011). We further discuss possible reasons for the different results among studies, particularly regarding learning and task requirements.

#### Outcome responses

Morris et al. (2004) and Apicella et al. (2011) examined TAN responses to probabilistic rewards. Morris et al. (2004) showed that TANs respond to probabilistic rewards without probability-based distinction of the responses during an instrumental conditioning task. Contrastingly, Apicella et al. (2011) showed that TAN responses to reward outcomes were dependent on the reward probability during a classical conditioning task. One possible reason for the discrepant results of these two studies is the different task structure for indicating the reward probability. Apicella et al. (2004) reported that monkeys did not observe a cue stimulus for the reward probability, and they needed to learn the reward probability in a block of trials, since the reward probability changed (0, 0.25, 0.75, or 1.0) suddenly every 40-70 trials. In the study by Morris et al. (2004), visual stimuli explicitly informed the monkeys of the reward probability (0, 0.25, 0.75, or 1.0) in each trial. The present results seem to be in line with findings by Morris et al. (2004). In our task, the reward probability was not explicitly cued to the monkeys but was predictably associated with the number of trials through a series of choices (Fig. 1E). It is possible that the monkeys tracked the reward probability, and thus the responses of TANs to the reward outcomes did not reflect the reward probability (Fig. 6).

In the present study, no responses to reward outcomes were observed during repeat trials, in which monkeys received rewards in almost 100% of the trials (Fig. 3A–F). This observation is generally consistent with the near absence of responses to the 100% rewards in the study by Apicella et al. (2011) (Fig. 4B in Apicella et al. 2011), but inconsistent with the clear responses to the 100%-reward condition in the study by Morris et al. (2004) (Fig. 4B in Morris et al., 2004). If the knowledge of reward probability determines the response modulations, the same results should have been observed in the study by Morris et al. (2004) and our study. Namely, these observations cannot be solely explained by knowledge of the reward probability. Another potential explanation is that the monkeys in our study may not have needed to learn which was the reward target during repeat trials, since the monkeys had already learned those relationships during the preceding search trials. In their study, monkeys chose the left or right lever and obtained rewards. Therefore, the monkeys may have learned which lever yielded rewards in every trial. As such, both the association between the stimulus and reward probability, and the association between behavioral responses and rewards may be key factors, which together are core features of thalamo-striatal signals (Yamanaka et al., 2018). Because TANs exhibit dynamic changes in the reward-related responses through learning (Aosaki et al., 1994b; Blazquez et al., 2002), the learning requirement is likely to have produced different TAN responses evoked by the probabilistic rewards among these three studies.

All three studies exhibited consistent results regarding the temporal patterns of the responses to reward and no-reward outcomes. Discharge increases were followed by decreases during reward outcomes (Figs. 3B and 4A, IF type; 4B in Morris et al., 2004; Fig. 4B in Apicella et al., 2011). In contrast, increases alone or activity decreases followed by increases were observed after no-reward outcomes (Figs. 3B, and 4D, F and SF types; 4B in Morris et al., 2004; Fig. 5B in Apicella et al., 2011). Although these two previous studies did not analyze the temporal patterns of feedback responses quantitatively, the responses seem to be consistent with our observations (Fig. 4). The distinct temporal patterns of responses may differentiate the acetylcholine release after reward and no-reward feedbacks.

#### Cue responses

Cue responses of TANs were consistently observed among the three studies. Morris et al. (2004) demonstrated similar TAN responses to the cue stimulus without reward probability-related response modulation (Fig. 4B in Morris et al., 2004). The study by Apicella et al. (2011) did not have a cue stimulus for reward probability, but TANs similarly responded to signals instructing the monkeys to move their arms, without probability modulation (Fig. 4B in Apicella et al., 2011), while TAN responses were sometimes observed for movement signals in other studies (Apicella et al., 1998; Yamada et al., 2004). Here, no cue stimulus for reward probability was used, but TANs responded consistently to the start cue, irrespective of trial type (Fig. 8). The observed temporal patterns of these responses were also consistent among the three studies, i.e., discharge decreases were occasionally followed by increases (Fig. 8B and C; Fig. 4B in Morris et al., 2004; Fig. 4B in Apicella et al., 2011).

### Adaptive behavior and reinforcement learning

Learning reward values from the consequences of selected choices is one of the core elements in the reinforcement learning model of the basal ganglia (Houk et al., 1995). Previous studies have shown that TANs detect and discriminate different kinds of motivational outcomes (Ravel et al., 2003; Yamada et al., 2004) and adapt their responses through learning (Aosaki et al., 1994b; Blazquez et al., 2002). In this study, TANs detected the reward and no-reward outcomes specifically during search trials, in which monkeys were required to learn target-reward associations for additional rewards.

Several recent studies have revealed that TANs are involved in various kinds of cognitive and motor performance during reward-based decision-making tasks (Atallah et al., 2014; Franklin and Frank, 2015; Nougaret and Ravel, 2015; Stalnaker et al., 2016). Some of these studies showed the specific roles of TANs in reinforcement learning, including signaling animal’s beliefs about the current state (Stalnaker et al., 2016) or signaling uncertainty, allowing the learning rate to be dynamically adjusted (Franklin and Frank, 2015). In line with the notion that the thalamo-striatal system controls acetylcholine signals for associative learning during environmental events (Yamanaka et al., 2018), the lack of TAN responses to reward feedback in repeat trials is likely due to the monkeys’ certainty regarding reward obtainment. Our findings suggest that TANs detect behavioral outcomes when they are not certainly predicted, i.e., TANs signal the outcomes, from which animals learn.

### Cholinergic signals in the striatal circuit

The striatum is involved in reinforcement learning and value-based choices under the influence of dopaminergic and cholinergic signals (Fig. 9). Acetylcholine signals conveyed by cholinergic interneurons, i.e., TANs, modulate the activity of striatal projection neurons directly and indirectly via fast-spiking inhibitory interneurons (Calabresi et al., 2000; Zhou et al., 2002). Indeed, presumed projection neurons, PANs, and presumed inhibitory interneurons, FSNs, demonstrate search trial-specific activity modulations like the feedback responses of TANs (Yamada et al., 2011; Yamada et al., 2016). Moreover, PAN and FSN activity is modulated by the target location and reward probability, while TAN activity is not. Thus, search-specific activity modulation in the striatum may originate from TANs, whereas target location and reward probability modulation may originate from other cell types or cortical inputs. The search-specific signals of TANs may be conveyed from the centromedian parafascicular (CM/Pf) complex in the intralaminar nuclei of the thalamus (Fig. 9, black arrows), which is suggested to control associability (Yamanaka et al., 2018). Dopamine neurons in the substantia nigra par compacta (SNs) send a reward prediction error signal to the dorsal striatum via their phasic responses to unpredicted reward outcomes during the multi-step choice task (Fig. 9, gray arrows) (Enomoto et al., 2011; Enomoto et al., 2020). This signal must modulate the striatal output activity (i.e., PANs), which signals an action or expected values of actions or search-repeat context (Yamada et al., 2011; Yamada et al., 2013). It is known that acetylcholine release in the striatum modulates dopamine release at the presynaptic terminal of the dopamine neurons via nicotinic acetylcholine receptors (Di Chiara et al., 1994; Jones et al., 2001; Zhou et al., 2001; Partridge et al., 2002; Cragg, 2006; Calabresi and Di Filippo, 2008; Surmeier et al., 2011). Thus, it is possible that cholinergic signals conveyed by TANs affect monkeys’ target valuation by modifying dopamine release in the local striatal circuit (Aosaki et al., 1994a; Schulz and Reynolds, 2013; Kim et al., 2019). Search-specific cholinergic signals may shape the striatal output activity directly and indirectly via modulating dopamine reward-prediction-error signals, suggesting that cholinergic signals in the striatum might control learning rate in reinforcement learning.

**Figure 9.**
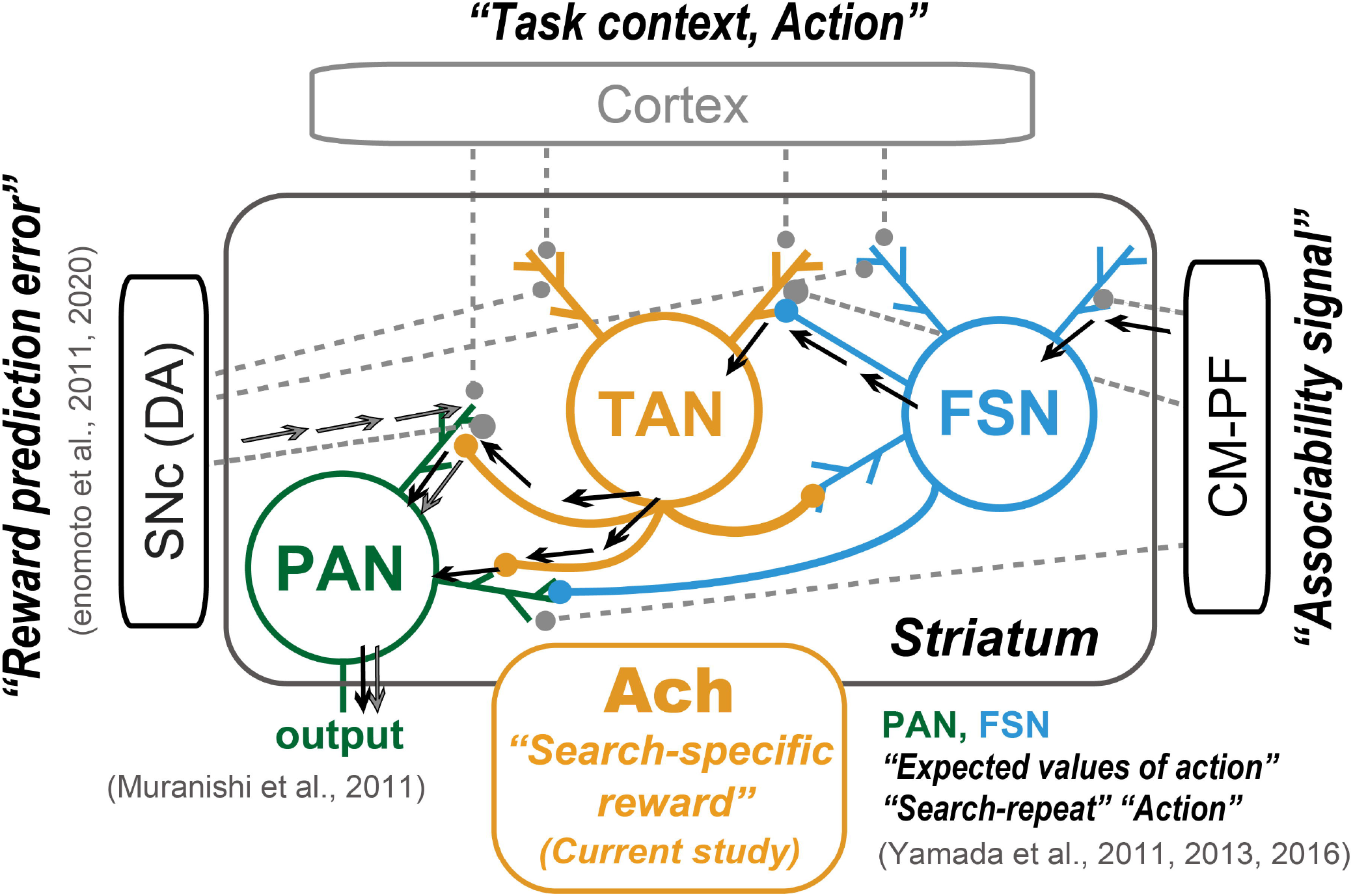
Role of TANs in the striatum circuit. Schematic drawing of the striatum circuit composed of TANs (presumed cholinergic interneurons, tonically active neuron), PANs (presumed output neurons, physically active neurons), and FSNs (presumed parvalbumin-containing GABAergic interneurons, fast-spiking neurons), which summarizes the signals of these neurons during the multi-step choice task. CM-PF, Centromedian (CM) and parafascicular (Pf) nuclei of the thalamus; SNc, substantia nigra par compacta. Ach; acetylcholine, DA: dopamine. The black arrows indicate two possible ways of how thalamic signals modulate the striatal output via pause of acetylcholine release. The gray arrows indicate how dopamine signals modulate the striatal output.

## Acknowledgments

The authors wish to express appreciation to Masayuki Matsumoto. The authors thank Yoshiko Yabana and Suwa Yuki for technical assistance.

## Author contributions

All authors designed the research. H.Y. and H.I. conducted the experiments. H.I. analyzed the data. All authors evaluated the analyzed results. H.Y. wrote the manuscript. H.I. wrote parts of the manuscript. All authors edited and approved the final manuscript.

